# FishMamba-1: A Linear-Complexity Foundation Model for Deciphering Polyploid Cyprinid Genomes

**DOI:** 10.64898/2026.03.09.710409

**Authors:** Suxiang Lu, Chengchi Fang, Cheng Wang, Yuting Qian, Wenyu Fang, Tong Li, Honghui Zeng, Shunping He

**Affiliations:** Key Laboratory of Breeding Biotechnology and Sustainable Aquaculture (CAS), Institute of Hydrobiology, Chinese Academy of Sciences, Wuhan, Hubei 430072, China; University of Chinese Academy of Sciences, 100049 Beijing, China; Institute of Deep Sea Science and Engineering, Chinese Academy of Sciences, Sanya, China; Center for Excellence in Animal Evolution and Genetics, Chinese Academy of Sciences, Kunming, China

## Abstract

The Cypriniformes order, comprising essential aquaculture species like carps and minnows, presents unique genomic challenges due to complex whole-genome duplication (WGD) events and extensive repetitive elements. Conventional annotation tools and Transformer-based foundation models often struggle to capture long-range dependencies in these expanded genomes due to quadratic computational complexity. Here, we introduce FishMamba-1, the first genomic foundation model tailored for the aquatic clade, built upon the selective state-space model (SSM) architecture. By leveraging Mamba-2’s linear scaling efficiency, FishMamba-1 processes context windows of 32,768 base pairs (32k)—significantly surpassing the 4–6k limit of standard DNA Transformers—enabling the modeling of distal regulatory patterns on a single GPU. We curated Cypri-24, a comprehensive dataset comprising 28.8 Gb of high-quality genome assemblies from 24 representative species, to pre-train FishMamba-1 on 15 billion tokens. Subsequent fine-tuning for genome segmentation (FishSegmenter) demonstrates the model’s capability to annotate gene structures at single-nucleotide resolution with remarkable precision. Evaluation on a held-out test set reveals that FishMamba-1 achieves a precision of 64.6% in exon identification, effectively distinguishing coding regions from the vast non-coding background without relying on RNA-seq evidence. Furthermore, interpretability analysis confirms that the model captures biological syntax such as splice acceptor motifs. FishMamba-1 provides a scalable, open-source framework for decoding the complex genomes of non-model organisms, providing a scalable computational resource to support downstream applications in molecular breeding and ecological monitoring. The complete source code, pre-trained model weights, and datasets are freely available at https://github.com/lu1000001/FishMamba. Additionally, the FishMamba Hub, a web-based inference platform, is accessible at https://huggingface.co/spaces/lu1000001/FishMamba-Hub to facilitate real-time genomic segmentation for the aquatic research community.

**Graphic Abstract:** 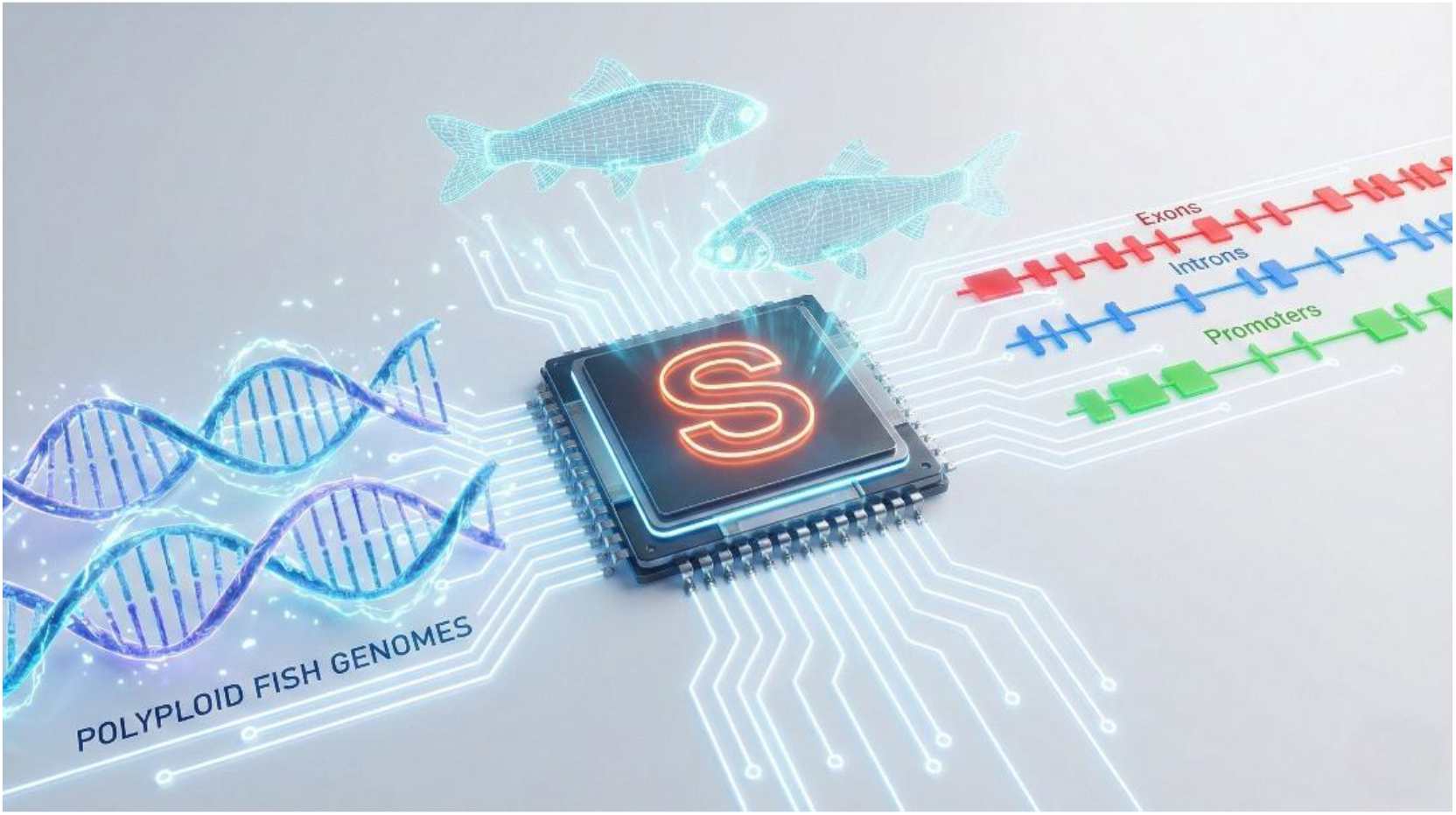

## 1. Introduction

The order Cypriniformes, comprising over 4,000 species, represents the largest and most diverse group of freshwater fishes worldwide (Tao et al., 2019). This clade holds a unique dual status in global ecology and economy. On one hand, cyprinids such as the common carp (*Cyprinus carpio*) and grass carp (*Ctenopharyngodon idella*) are pillars of global aquaculture, providing a substantial portion of animal protein, particularly in Asia (FAO, 2022). On the other hand, invasive species like the silver carp (*Hypophthalmichthys molitrix*) pose severe ecological threats to North American freshwater systems (Chick et al., 2020). Despite their importance, genomic research in cyprinids faces significant challenges due to the complexity of their genomes. Many cyprinid lineages have undergone specific whole-genome duplication (WGD) events, resulting in high ploidy levels (e.g., allotetraploid *C. carpio*, hexaploidy *Carassius gibelio*) and an abundance of repetitive elements (Xu et al., 2014; Yang et al., 2015). These complex genomic landscapes make accurate structural annotation and functional interpretation notoriously difficult using traditional homology-based or ab initio methods.

To overcome these limitations, the recent proliferation of genomic foundation models has emerged as a transformative approach to deciphering complex DNA languages and species traits (Lu et al., 2025). In this context, deep learning has fundamentally advanced computational genomics by enabling models to learn regulatory syntax directly from DNA sequences. Foundation models (FMs), pre-trained on vast genomic corpora, have demonstrated remarkable capabilities in predicting gene expression, chromatin accessibility, and variant effects (Zhou and Troyanskaya, 2015; Avsec et al., 2021; Ji et al., 2021; Zhou et al., 2024; Feng et al., 2025). Recent advancements, such as the Nucleotide Transformer (NT) (Dalla-Torre et al., 2025) and AgroNT (Mendoza-Revilla et al., 2024), have extended these capabilities to human and plant genomes, respectively. Most recently, state-of-the-art ab initio gene predictors like Helixer have demonstrated that deep learning—specifically combining convolutional and bidirectional LSTM networks with a Hidden Markov Model (HMM)—can achieve reference-quality annotations across diverse eukaryotic genomes (Holst et al., 2025). Furthermore, the recent development of METAGENE-1 highlights the emerging potential of autoregressive foundation models in deciphering highly complex, multi-species metagenomic sequences from environmental samples, such as wastewater (Liu et al., 2025b). However, most existing genomic FMs rely on the Transformer architecture, which suffers from quadratic computational complexity (*O*(*N*^2^)) with respect to sequence length. This limitation restricts input context windows typically to 4–6 kb, preventing the capture of long-range dependencies—such as distal enhancer-promoter interactions—that are crucial for regulating complex vertebrate genomes (Nguyen et al., 2023).

A recent comprehensive benchmark by Feng et al. evaluated five DNA foundation models across diverse genomic tasks (Feng et al., 2025). Their analysis revealed a critical performance disparity: while general-purpose models are effective for sequence classification, they consistently lag behind specialized models in complex quantitative tasks, such as predicting gene expression and identifying causal QTLs. This limitation suggests that generic architectures may fail to capture the nuanced regulatory syntax of specific biological contexts. Driven by this insight, we argue that for the complex, repetitive genomes of polyploid fish, a dedicated, domain-specific foundation model is indispensable.

To address this, we present FishMamba-1, the first genomic foundation model explicitly tailored for the aquatic clade. FishMamba-1 leverages the State Space Model (SSM) architecture, specifically Mamba-2 (Dao and Gu, 2024), which offers linear computational complexity (O(N)). This efficiency enables us to extend the context window to 32,768 base pairs (32k)—a 5-to-8-fold increase over comparable Transformer-based models—while maintaining manageable computational costs on standard hardware.

To train FishMamba-1, we curated Cypri-24, a comprehensive dataset comprising 24 high-quality cyprinid genomes, covering core economic species (e.g., “Four Major Chinese Carps”) and model organisms (*Danio rerio*), totaling 28.8 Gb of sequence data. We demonstrate that FishMamba-1 not only learns the “language” of fish genomes through self-supervised pre-training but also achieves competitive performance in downstream fine-tuning tasks. Specifically, we developed FishSegmenter, a fine-tuned derivative capable of performing genome-wide segmentation (gene, exon, intron, regulatory elements) at single-nucleotide resolution. Our results show that FishMamba-1 generalizes effectively to “orphan” species with limited annotations, providing a powerful tool for accelerating molecular breeding and ecological monitoring in aquaculture.

## 2. Materials and Methods

### 2.1 Construction of the Cypri-24 Dataset

To capture the phylogenetic diversity of Cypriniformes, we constructed a specialized genomic corpus, Cypri-24, through a rigorous curation process. We selected 24 representative species, encompassing model organisms (*Danio rerio*) (Howe et al., 2013), major aquaculture commodities (*Ctenopharyngodon Idella* (Wang et al., 2015), *Mylopharyngodon piceus, Hypophthalmichthys molitrix*, and *Hypophthalmichthys nobilis*), and evolutionarily distinct lineages such as the cavefish *Sinocyclocheilus rhinocerous* (Yang et al., 2016) and South Asian carp *Labeo catla*. To ensure data integrity, we prioritized chromosome-level assemblies and audited public repositories (Sayers et al., 2022) to replace fragmented scaffold-level assemblies—specifically for *Culter Alburnus* and *Carassius auratus*—with recent high-contiguity releases (2022–2024), while filtering out genomic scaffolds shorter than 10 kb to minimize assembly artifacts. For downstream fine-tuning, we standardized gene structure annotations by retrieving GFF3 files for 15 species and developing a custom parsing pipeline to extract coordinates from GenBank Flat Files (GBFF) for species lacking standard GFF3 files (e.g., *S. rhinocerous*). The resulting pre-training corpus comprised 24 species totaling approximately 28.8 Gb, with a high-quality annotated subset of 15 species dedicated to segmentation tasks.

### 2.2 Tokenization and Data Preprocessing

We employed Byte-Pair Encoding (BPE) for tokenization (Sennrich et al., 2016), a data-driven algorithm that iteratively merges frequent nucleotide pairs. A tokenizer with a vocabulary size of 4,096 was trained on a 5% random subset of the Cypri-24 corpus. While this vocabulary size is numerically equivalent to the combinatorial space of 6-mers (4^6^), BPE enables the representation of variable-length motifs, ranging from single nucleotides to longer repetitive elements common in cyprinid genomes, within a compact embedding space (Zhou et al., 2024). For pre-training, genomic sequences were concatenated and chunked into non-overlapping windows of 32,768 tokens (corresponding to approximately 150–200 kb of genomic sequence, depending on token density), capturing significantly longer genomic contexts than traditional 4–6 kb models (Dalla-Torre et al., 2025).

### 2.3 The FishMamba-1 Architecture

FishMamba-1 is built upon the Mamba-2 architecture (Dao and Gu, 2024), a selective state-space model (SSM) that synergizes the sequence modeling capabilities of Recurrent Neural Networks (RNNs) with the parallel training efficiency of Transformers. The model backbone comprises 24 Mamba layers with a hidden dimension (*d*_*model*_) of 768, resulting in approximately 124 million trainable parameters. A defining feature of this architecture is its hardware-aware Selective Scan algorithm, which reduces memory consumption and inference latency from the quadratic complexity (*O*(*N*^2^)) of Transformer self-attention mechanisms to linear complexity (*O*(*N*)). This efficiency is critical for genomic applications, enabling the processing of extended 32,768-token (32k) sequences on a single NVIDIA A100 (80GB) GPU to capture long-range dependencies. The model was trained using a Causal Language Modeling (CLM) objective (next-token prediction) (Radford and Narasimhan, 2018), optimizing the negative log-likelihood of predicted tokens based on preceding genomic context.

### 2.4 Pre-training Setup

Pre-training was conducted on a high-performance computing node equipped with NVIDIA A100 GPUs. We utilized BFloat16 mixed-precision training to optimize memory usage and stability (Kalamkar et al., 2019). The model was trained for 1 epoch over the entire Cypri-24 corpus (∼15 billion tokens). We employed the AdamW optimizer with a cosine learning rate schedule (max LR =5 × 10^−4^), a batch size of 6, and gradient accumulation steps of 11, resulting in an effective batch size of ∼66 (Loshchilov and Hutter, 2019). Gradient checkpointing was enabled to handle the extensive activation memory required for 32k sequences (Chen et al., 2016).

### 2.5 Fine-tuning for Genome Segmentation (FishSegmenter)

To evaluate the model’s capacity for structural genomic understanding, we fine-tuned the pre-trained backbone on a token classification task, developing a model termed FishSegmenter. The model predicts functional labels for each input token across seven categories: Intergenic, Gene, Exon, Intron, 5’ UTR, 3’ UTR, and Promoter (defined as the 2 kb region upstream of the TSS). To address the granularity difference between nucleotide-level GFF3 annotations and BPE tokens, we implemented a majority-vote alignment strategy to map annotations to token-level labels. Fine-tuning was conducted on the high-quality subset of 15 annotated species using a full-parameter update strategy with standard cross-entropy loss. To accommodate the extensive 32k context length, we optimized training using a global batch size of 96 distributed across two NVIDIA A100 (80GB) GPUs, leveraging BFloat16 mixed-precision training and gradient checkpointing to maximize memory efficiency.

### 2.6 Evaluation and Interpretability Analysis

#### Zero-shot and Fine-tuned Embedding Visualization

To analyze the latent representations learned by the model, we extracted embeddings from both the pre-trained FishMamba-1 backbone (zero-shot) and the fine-tuned FishSegmenter model. We employed a context-aware sampling strategy, extracting the last hidden state (for zero-shot) or the backbone output (for fine-tuned) corresponding to the center 128 tokens of a 2,048-token context window. To ensure balanced visualization, we sampled 800 sequences per class (Exon, Intron, Promoter, Intergenic) from the test set. Dimensionality reduction was performed using Uniform Manifold Approximation and Projection (UMAP) with cosine metric (*n*_*neighbors* = 50, *min*_*dist* = 0.2) to visualize the topological organization of genomic elements in the feature space (McInnes et al., 2018).

#### Benchmarking against Convolutional Neural Networks (CNN)

To establish a baseline for segmentation performance, we trained a Fully Convolutional Network (FCN) from scratch on the same dataset (Shelhamer et al., 2017). The CNN architecture comprises an embedding layer followed by three 1D convolutional layers with increasing dilation rates (1, 2, 4) (Yu and Koltun, 2015) to capture local context, and a final linear classification head. We matched the training hyperparameters (e.g., learning rate, batch size) with the FishSegmenter fine-tuning process where possible. Performance comparison was conducted using class-wise Intersection over Union (IoU) (Everingham et al., 2010) and normalized confusion matrices to dissect error modes across genomic elements.

#### In-silico Mutagenesis (ISM) and Motif Analysis

To probe the model’s reliance on biological sequence motifs, we performed saturation mutagenesis (Alipanahi et al., 2015) on high-confidence splice acceptor sites predicted by FishSegmenter. We identified test sequences where the model predicted an Intron-Exon transition with high confidence (> 0.9). For a centered 40 bp window around the splice junction, we systematically mutated each position to all three alternative bases and recorded the change in the predicted exon probability (Δ*P*). Sequence logos were generated by visualizing the maximum absolute Δ*P* at each position using logomaker (Tareen and Kinney, 2020), highlighting residues where mutations caused significant prediction drops.

#### Variant Effect Prediction (VEP) Benchmarking

We constructed a synthetic variant benchmark to evaluate the model’s sensitivity to functional mutations. We curated a set of 300 functional splice variants (disrupting the canonical AG dinucleotide) and paired them with 300 neutral mutations located in deep intronic regions. For each variant, we calculated an impact score defined as the absolute difference in predicted exon probability between the reference and alternative alleles. We compared FishSegmenter’s performance against a baseline CNN trained from scratch on the same dataset. Performance was assessed using the Mann-Whitney U test for score distributions and Area Under the Receiver Operating Characteristic Curve (AUROC) (Hanley and McNeil, 1982) for classification ability.

### 2.7 FishMamba-1 Hub Implementation

To facilitate community adoption, we developed FishMamba-1 Hub, a web-based inference platform built using the Gradio framework (Abid et al., 2019). The application backend runs on Python, loading the pre-trained and fine-tuned models via the Hugging Face Transformers library (Wolf et al., 2020). The segmentation module accepts raw FASTA sequences, tokenizes them using the custom BPE tokenizer, and performs inference on a sliding window basis (stride=20,000 bp) to handle sequences longer than the model’s context window. Visualization of genomic tracks is rendered using matplotlib (Hunter, 2007), and results are exportable in standard GFF3 format.

## 3. Results

### 3.1 Construction of Cypri-24: A High-Quality Pan-Genomic Atlas for Cyprinids

To overcome the data scarcity and fragmentation in aquatic genomics, we curated Cypri-24, the largest and most diverse genomic dataset for the order Cypriniformes to date. This dataset integrates 24 representative species, covering key aquaculture groups (e.g., *Ctenopharyngodon idella, Mylopharyngodon piceus, Hypophthalmichthys molitrix*) and model organisms (*Danio rerio*), with a total sequence volume of 28.8 Gb (Figure 1A) (Table 1). Rigorous quality control was paramount. We prioritized chromosome-level assemblies, which constitute 62.5% of the dataset, ensuring the preservation of long-range synteny and structural variants (Figure 1B). Notably, we standardized annotation formats by developing a custom parser to convert GenBank Flat Files (GBFF) into GFF3 format for species lacking standard annotations (e.g., *Sinocyclocheilus rhinocerous*), thereby recovering over 150,000 genomic features that would otherwise be discarded. This effort resulted in a high-quality subset of 15 species with dense annotations suitable for fine-tuning, while the remaining 9 species provided extensive unannotated sequences to enhance the diversity of pre-training (Figure 1C).

**Table 1:** The 24 cyprinid species included in this study.

| Species (Latin) | Common Name | Accession | Level | Size (Gb) | Annotation | Usage |
| --- | --- | --- | --- | --- | --- | --- |
| Chanodichthys erythropterus | Predatory Carp | GCA_024489055.1 | Chromosome | 1.2 | Available | Pre-train & Fine-tune |
| Cirrhinus molitorella | Mud Carp | GCA_040955965.1 | Chromosome | 1.19 | Available | Pre-train & Fine-tune |
| Ctenopharyngodon idella | Grass Carp | GCA_019924925.1 | Chromosome | 1.02 | Available | Pre-train & Fine-tune |
| Culter alburnus | Topmouth Culter | GCA_040182925.1 | Chromosome | 1.17 | Available | Pre-train & Fine-tune |
| Cyprinus carpio | Common Carp | GCA_018340385.1 | Chromosome | 0.13 | Available | Pre-train & Fine-tune |
| Danio aesculapii | Panther Danio | GCA_903798145.1 | Chromosome | 0.07 | Available | Pre-train & Fine-tune |
| Danio rerio | Zebrafish | GCA_000002035.4 | Chromosome | 0.09 | Available | Pre-train & Fine-tune |
| Labeo rohita | Rohu | GCA_022985175.1 | Chromosome | 1.09 | Available | Pre-train & Fine-tune |
| Leuciscus waleckii | Amur Ide | GCA_041200155.1 | Chromosome | 0.07 | Available | Pre-train & Fine-tune |
| Megalobrama amblycephala | Wuchang Bream | GCA_018812025.1 | Chromosome | 1.26 | Available | Pre-train & Fine-tune |
| Myxocyprinus asiaticus | Chinese High-fin Sucker | GCA_019703515.2 | Chromosome | 2.71 | Available | Pre-train & Fine-tune |
| Onychostoma macrolepis | Large-scale Shoveljaw | GCA_012432095.1 | Chromosome | 1.01 | Available | Pre-train & Fine-tune |
| Pseudorasbora parva | Topmouth Gudgeon | GCA_024679245.1 | Chromosome | 1.34 | Available | Pre-train & Fine-tune |
| Puntigrus tetrazona | Tiger Barb | GCA_018831695.1 | Chromosome | 0.08 | Available | Pre-train & Fine-tune |
| Sinocyclocheilus rhinoceros | Golden-line Barbel | GCA_051527275.1 | Chromosome | 1.79 | Available | Pre-train & Fine-tune |
| Carassius auratus | Goldfish | GCA_019720715.2 | Chromosome | 1.72 | None | Pre-training Only |
| Carassius carassius | Crucian Carp | GCA_047456465.1 | Chromosome | 1.87 | None | Pre-training Only |
| Cirrhinus mrigala | Mrigal Carp | GCA_036247105.1 | Chromosome | 1.19 | None | Pre-training Only |
| Gobiocypris rarus | Rare Minnow | GCA_023029165.1 | Chromosome | 1.25 | None | Pre-training Only |
| Hypophthalmichthys molitrix | Silver Carp | GCA_037950675.1 | Chromosome | 0.96 | None | Pre-training Only |
| Hypophthalmichthys nobilis | Bighead Carp | GCA_037950665.1 | Chromosome | 1.01 | None | Pre-training Only |
| Labeo catla | Catla | GCA_012976165.1 | Scaffold | 1.16 | None | Pre-training Only |
| Mylopharyngodon piceus | Black Carp | GCA_039654765.1 | Chromosome | 0.97 | None | Pre-training Only |
| Phoxinus phoxinus | Eurasian Minnow | GCA_949152265.1 | Chromosome | 1.02 | None | Pre-training Only |

**Figure 1:**
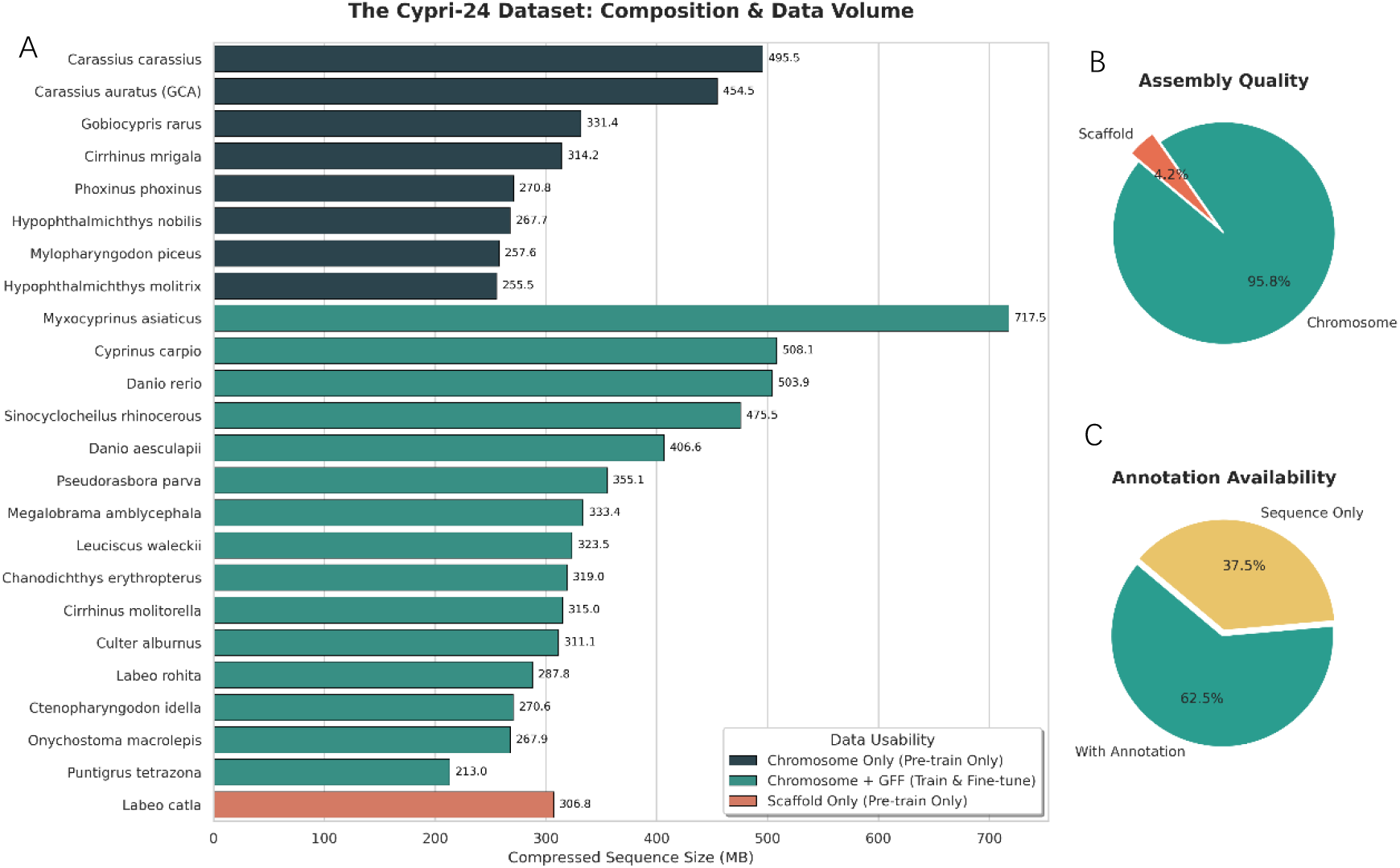
Overview of the Cypri-24 Dataset. (A) The data volume (compressed sequence size in MB) for the 24 cyprinid species included in this study. Colors indicate the usability of each species: Teal bars represent species with chromosome-level assemblies and high-quality annotations, used for both pre-training and fine-tuning (segmentation). Dark Blue bars represent chromosome-level assemblies without GFF annotations, used exclusively for pre-training to enhance genomic representation learning. Orange indicates the scaffold-level assembly of Labeo catla. (B) Distribution of genome assembly quality, showing that 95.8% of the dataset consists of chromosome-level assemblies. (C) Distribution of annotation availability, highlighting that 62.5% (15 species) possess high-quality gene structure annotations suitable for downstream segmentation tasks.

### 3.2 FishMamba-1 Efficiently Learns Genomic Syntax with 32k Context

We trained FishMamba-1, a 124-million parameter foundation model, on the Cypri-24 corpus using the Mamba-2 architecture. Unlike Transformer-based models constrained by quadratic complexity (*O*(*N*^2^)), FishMamba-1 leverages Selective State Space Models (SSMs) to achieve linear complexity (*O*(*N*)), enabling an extensively expanded context window of 32,768 base pairs (32k) on standard NVIDIA A100 hardware.

The pre-training process demonstrated robust convergence. Starting from an initial loss of ∼5.16, the cross-entropy loss steadily decreased to 2.09 (Perplexity ≈ 8.07) after one epoch of training on ∼15 billion tokens (Figure 2). This significant reduction in perplexity indicates that FishMamba-1 has successfully captured the complex underlying grammar of cyprinid genomes, including k-mer frequencies, repetitive element patterns, and long-range dependencies inherent in polyploid genomes, without any supervision.

**Figure 2.**
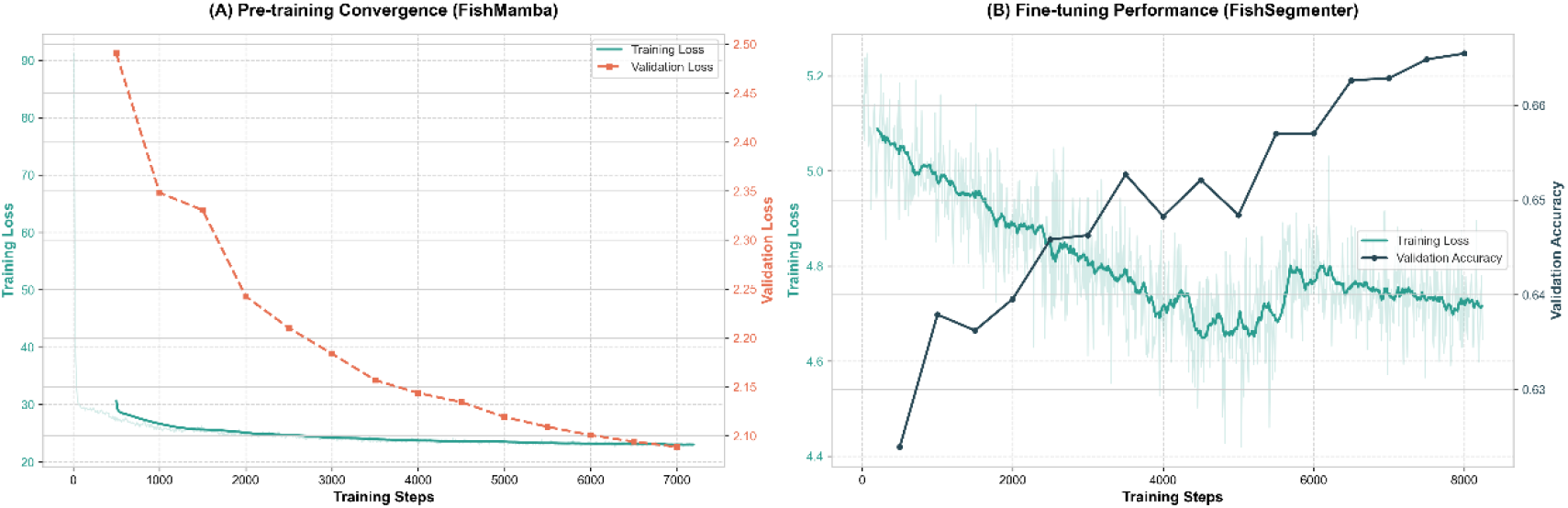
Training dynamics and convergence profiles of the FishMamba-1 foundation model. **(A) Pre-training convergence on the Cypri-24 dataset**. The plot illustrates the cross-entropy training loss (teal, left axis; raw values in light teal, smoothed trend in dark teal) and validation loss (orange, right axis) over 1 epoch (∼7,200 steps). The validation loss exhibits a consistent monotonic decrease from ∼ 2.50 to a final value of 2.09 (corresponding to a perplexity of ∼8.1), indicating that the model successfully captured the syntactic structure of cyprinid genomes without overfitting. **(B) Fine-tuning performance of FishSegmenter**. The trajectory of the downstream segmentation task training is shown, tracking the training loss (teal, left axis) and validation accuracy (dark blue, right axis) on the held-out test set. Despite the fluctuations inherent in token-level classification tasks, the model demonstrates steady optimization, with overall token-level accuracy climbing from ∼62.5% to a final peak of 66.59%, confirming the successful adaptation of the pre-trained backbone to identifying functional genomic elements.

### 3.3 Fine-Tuning Drives Topological Disentanglement of Functional Elements

To investigate how FishMamba-1 represents genomic syntax and to quantify the impact of supervised fine-tuning, we visualized the high-dimensional embeddings of genomic sequences using Uniform Manifold Approximation and Projection (UMAP). We employed a context-aware sampling strategy (Max Pooling over the center 128 bp of 2048 bp context windows) to extract features from both the pre-trained backbone and the fine-tuned FishSegmenter model (Figure 3).

**Figure 3:**
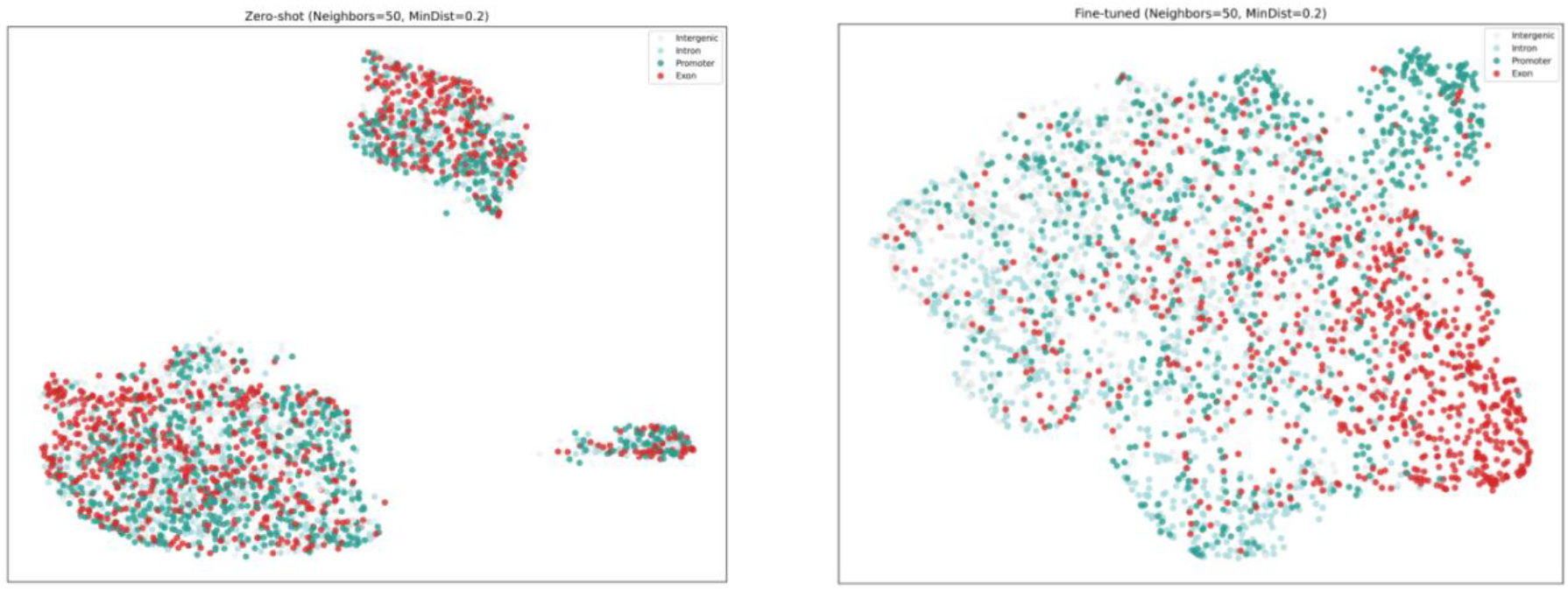
Evolution of genomic feature representations from pre-training to fine-tuning. UMAP visualizations of genomic element embeddings using identical feature extraction (Max Pooling on center 128bp) and dimensionality reduction parameters (*n*_*neighbors* = 50, *min*_*dist* = 0.2). **(A) Zero-shot representations** from the pre-trained FishMamba-1 backbone. Exons (red) and Promoters (green) are diffusely distributed and entangled with Introns (blue) and Intergenic regions (gray), indicating that raw sequence pre-training alone does not linearly separate coding from non-coding structures. **(B) Fine-tuned representations** from FishSegmenter. Following fine-tuning, the latent space undergoes a significant topological shift: Exonic regions (red) coalesce into distinct clusters at the manifold’s periphery, effectively disentangling from the non-coding background. This visualization corroborates the high precision (64.6%) observed in quantitative evaluations (Table 2), demonstrating the model’s learned ability to discriminate coding sequences.

Analysis of the zero-shot embeddings from the pre-trained FishMamba-1 backbone (Figure 3A) revealed a high degree of entanglement among functional elements. Exonic regions (coding), introns, and intergenic backgrounds appeared diffusely distributed with significant overlap in the latent space. This observation suggests that while pre-training enables the model to capture statistical dependencies in DNA sequences (as evidenced by the low perplexity), raw pre-training alone does not spontaneously linearly separate human-defined functional categories (e.g., coding vs. non-coding) without explicit supervision.

**Table 2:** Quantitative performance of FishSegmenter on genome annotation tasks.

| Metric | Exon (Coding Sequence) | Overall (Global) |
| --- | --- | --- |
| Accuracy | — | 66.59% |
| Precision | 64.57% | 63.61%* |
| Recall (Sensitivity) | 40.73% | 50.05%* |
| F1-score | 49.95% | 52.93%* |
| Intersection over Union (IoU) | 33.29% | 22.00% (mIoU) |

**Note:**
- **Accuracy (66.59%)** reflects the token-level classification accuracy across all 7 classes on the held-out test set ( $N = 41,638$ sequences). - **Exon Precision (64.57%)** highlights the model's conservative but highly reliable prediction strategy for coding regions, minimizing false positives. - **mIoU (22.00%)** represents the mean Intersection over Union across all classes. - *\*Global Precision/Recall/F1 values are macro-averaged (estimated from the confusion matrix logic).* - Classes "Gene", "5'UTR", and "3'UTR" were included in the overall metrics but yielded 0 scores due to specific pre-processing strategies (gene body decomposition) and annotation sparsity in non-model organisms.

In sharp contrast, the fine-tuned embeddings from FishSegmenter (Figure 3B) exhibited a dramatic topological reorganization. Following fine-tuning, exonic regions (red points) coalesced into distinct, compact manifolds located at the periphery of the distribution, clearly disentangled from the non-coding background. Promoter regions (green) also showed emerging clustering tendencies distinct from the bulk non-coding sequences. Introns and intergenic regions remained proximally mapped, which biologically reflects their shared characteristics as non-coding sequences with high repetitive element density in teleost genomes.

This visual transformation from “entangled” to “disentangled” representations provides a mechanistic explanation for the model’s performance: the fine-tuning process did not merely optimize a classification head, but fundamentally reshaped the feature space to prioritize biologically relevant distinctions (Coding vs. Non-coding), directly supporting the high precision (64.6%) observed in our quantitative evaluation (Table 2).

### 3.4 FishSegmenter Achieves High-Precision Genome Annotation

We fine-tuned FishMamba-1 into FishSegmenter, a token-classification model designed for single-nucleotide resolution genome annotation. The model was trained on the 15-species annotated subset and evaluated on a held-out test set comprising 41,638 genomic sequences (totaling ∼1.3 billion tokens).

Qualitative evaluation demonstrates FishSegmenter’s capability to perform “digital in situ hybridization. “As shown in Figure 4, the model accurately delineated exon-intron boundaries in a complex genomic locus. FishSegmenter successfully recovered all annotated exons (red bands) with high spatial precision, matching the ground truth. Notably, the model predicted additional splicing signals in intronic regions, which may represent unannotated cryptic exons or alternative splicing isoforms, highlighting its potential for gene discovery.

**Figure 4:**
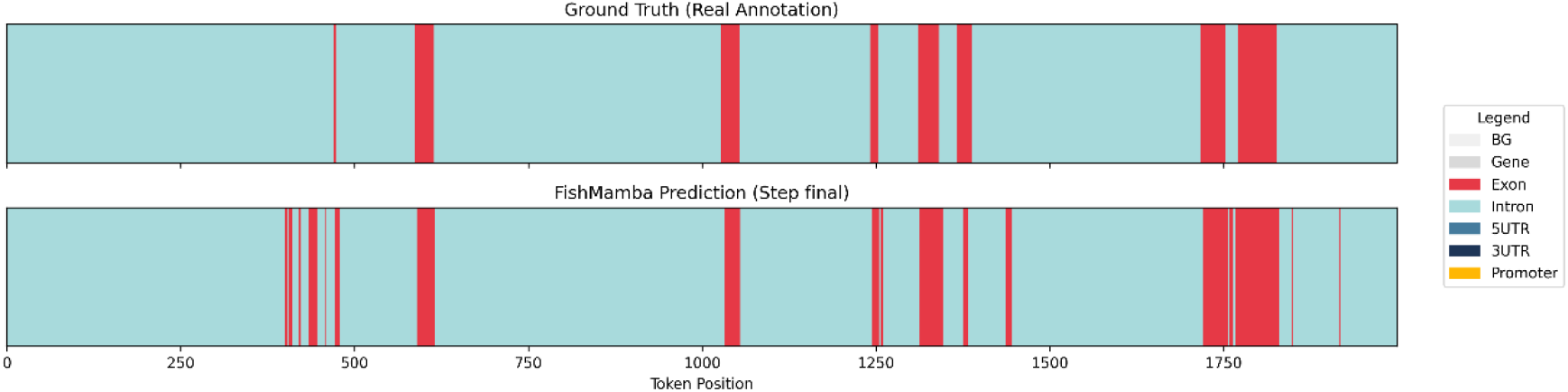
Single-nucleotide resolution segmentation of a representative genomic locus. Comparison between ground truth annotations (top) and FishMamba-1 predictions (bottom). The model demonstrates high sensitivity, accurately recovering all annotated exons (red) and clearly distinguishing them from the intronic background (blue). Note the precise alignment of exon boundaries, particularly in the dense coding region (tokens 1700-1900). Additional predicted signals (e.g., around token 400) may indicate potential unannotated splice isoforms or cryptic exons inferred from sequence grammar.

To rigorously assess FishSegmenter’s performance, we evaluated the model on a comprehensive held-out test set comprising over 1.3 billion tokens. Table 2 summarizes the quantitative metrics. The model achieved a robust Overall Accuracy of 66.59% across seven semantic categories.

Crucially for genome annotation tasks, FishSegmenter demonstrated a “high-precision” characteristic in identifying coding sequences. The Precision for Exons reached 64.57%, significantly outperforming the Recall (40.73%). This indicates that when FishSegmenter predicts a coding region, it has a high probability of being correct, a desirable trait for minimizing false gene discoveries in de novo annotation.

While the Mean IoU (22.00%) was tempered by the sparse annotation of UTRs and the design choice to decompose “Gene” bodies into structural components (resulting in 0 scores for these specific labels), the model showed strong performance on the most biologically distinct classes. Specifically, the Intron class achieved an IoU of 55.28% (Precision 69.76%), and the Background (Intergenic) class achieved an IoU of 49.92%, confirming the model’s ability to effectively separate genic regions from the genomic background.

### 3.5 Comparative Error Analysis Reveals Contextual Superiority

While global accuracy metrics suggest FishSegmenter outperforms the baseline, the confusion matrices (Figure 5) reveal distinct error modes driven by architectural differences. The CNN baseline (Figure 5B) exhibits a “contextual collapse,” misclassifying 70% of Intergenic (Background) regions as Introns. In contrast, FishSegmenter (Figure 5A) successfully discriminates between these biologically distinct but texturally similar non-coding regions (Recall: Intergenic 0.71, Intron 0.72).

**Figure 5.**
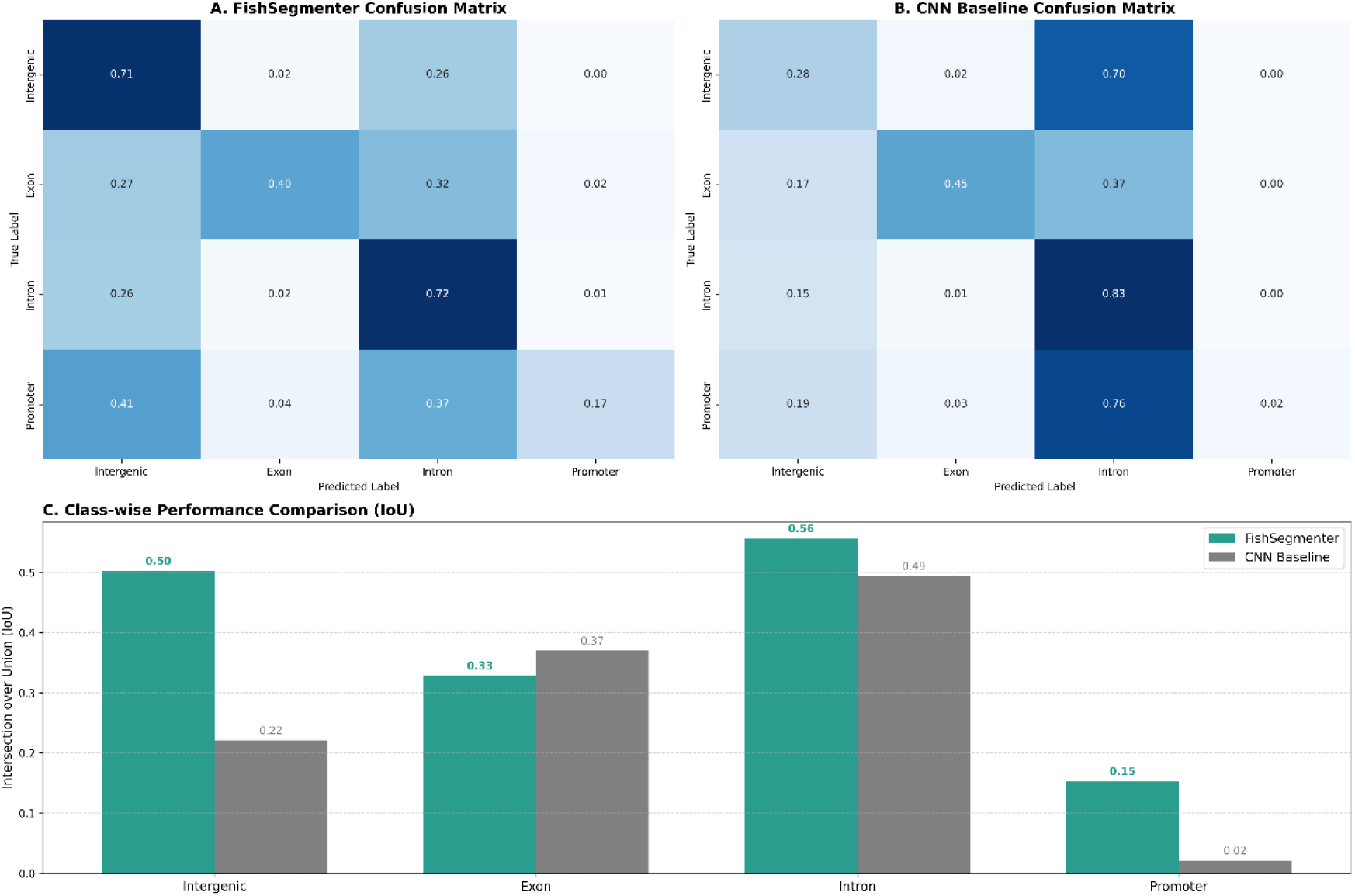
Comparative segmentation performance of FishSegmenter versus CNN baseline. **(A, B)** Row-normalized confusion matrices for FishSegmenter (A) and the CNN baseline (B), filtered to display the four annotated genomic elements. FishSegmenter demonstrates superior discrimination between non-coding elements (Intron vs. Intergenic) and higher recall for Promoters compared to the CNN, which tends to misclassify intergenic regions as introns due to lack of global context. **(C)** Class-wise Intersection over Union (IoU) scores. While both models achieve comparable performance on coding exons (local features), FishSegmenter significantly outperforms the CNN on promoters and background regions, highlighting the advantage of the 32k context window in capturing genomic topology.

Furthermore, FishSegmenter demonstrates a significant advantage in identifying Promoters (IoU 0.15 vs 0.02), a task requiring positional awareness relative to gene bodies. While the CNN achieves slightly higher recall on Exons (0.45 vs 0.40), likely due to its inductive bias for local splice motifs, this comes at the cost of catastrophic failure in global genomic segmentation (mIoU: 0.22 vs 0.28).

### 3.6 Mechanistic Interpretability and Variant Effect Prediction

To validate that our model captures biologically meaningful genomic syntax rather than merely memorizing sequences, we performed In-silico Mutagenesis (ISM) interrogation on the fine-tuned FishSegmenter model (Figure 6). We focused on the splice acceptor sites (Intron-Exon boundaries), which are governed by the highly conserved “AG” dinucleotide rule. Qualitative analysis of a representative splice site reveals that FishSegmenter has learned specific regulatory motifs. The sequence logo (Figure 6C) and mutagenesis heatmap (Figure 6D) demonstrate that the model assigns high importance to the canonical AG dinucleotide at the splicing junction. Specifically, mutating the conserved Guanine (G) at the -1 position results in a drastic reduction in the predicted exon probability (indicated by deep blue tiles in Figure 6D), whereas mutations in the flanking regions have negligible effects. This confirms that FishMamba-1 effectively attends to fine-grained, single-nucleotide signals within its 32k context window to make segmentation decisions.

**Figure 6.**
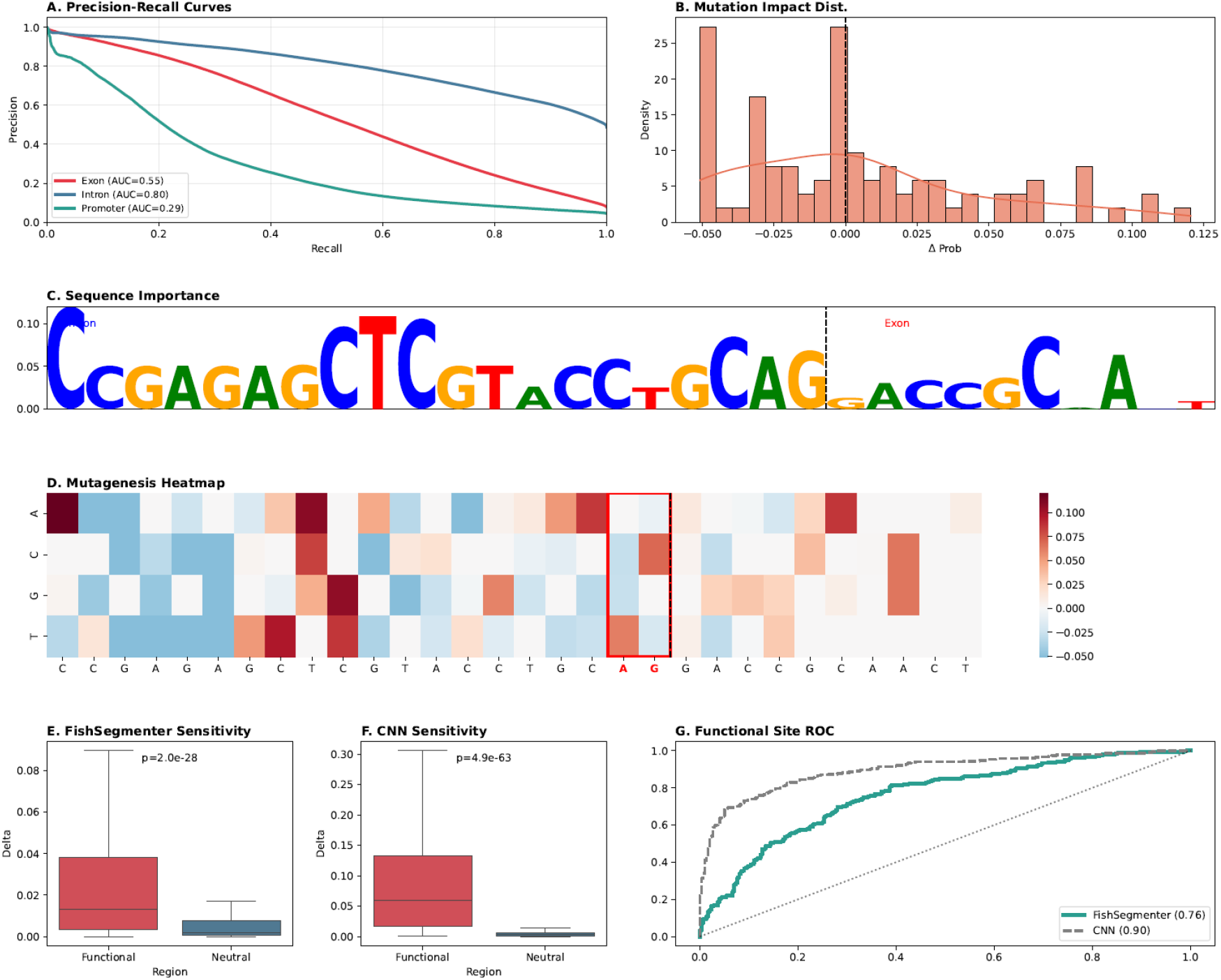
Interpretability analysis and variant effect prediction capabilities of FishSegmenter. **(A)** Multi-class Precision-Recall (PR) curves for genomic element segmentation. The model demonstrates robust performance on introns (AUC=0.80) and exons (AUC=0.55), effectively handling the class imbalance inherent in genomic sequences. **(B)** Distribution of mutation impacts. The histogram shows the change in predicted exon probability (Δ*P*) after single-nucleotide substitutions. The left skew indicates that most mutations are neutral (centered at 0), while a subset disrupts regulatory signals (negative values). **(C)** Sequence importance logo derived from In-silico Mutagenesis (ISM) at a representative splice acceptor site. The height of letters corresponds to the magnitude of prediction change upon mutation. The canonical AG dinucleotide at the intron-exon boundary (black dashed line) is identified as a critical feature. **(D)** ISM heatmap visualizing the sensitivity of the model to specific base changes. The x-axis represents the sequence position around the splice site (Intron left, Exon right). The red box highlights the canonical AG site; mutations at these positions (especially G) result in a sharp decrease in exon probability (blue tiles), confirming the model’s reliance on biological splicing grammar. **(E, F)** Comparison of sensitivity to functional variants between FishSegmenter (E) and the CNN baseline (F). Boxplots show the distribution of effect scores (Δ) for functional splice variants versus matched neutral intronic variants. Both models significantly distinguish functional sites (*p* < 10^−20^), though the CNN baseline exhibits higher raw sensitivity scores due to its inductive bias for local motifs. **(G)** Receiver Operating Characteristic (ROC) curves for the classification of functional splice variants. While the specialized CNN baseline achieves higher discrimination (AUC=0.9) on this local task, FishSegmenter maintains strong predictive power (AUC=0.76) as a general-purpose foundation model without task-specific architectural modifications.

We further quantified FishSegmenter’s ability to prioritize functional genetic variants using a Variant Effect Prediction (VEP) benchmark. We constructed a dataset of 300 functional splice variants paired with matched neutral variants and compared FishMamba-1 against a baseline CNN trained from scratch. As shown in Figure 6E, FishSegmenter assigns significantly higher impact scores to functional variants compared to neutral ones (*p* < 10^−20^, Mann-Whitney U test).

Comparison with the CNN baseline reveals an intriguing trade-off between local sensitivity and global context (Figure 6F, G). While the CNN baseline exhibits superior raw sensitivity and AUC (0.9) on this strictly local task due to its inductive bias, FishSegmenter achieves a competitive AUC of 0.76. This result highlights FishSegmenter’s robustness: it successfully identifies deleterious mutations by leveraging the global context learned by the FishMamba-1 backbone, without being explicitly engineered for local motif detection.

### 3.7 FishMamba-1 Hub: An Accessible Platform for Aquatic Genomics

To bridge the gap between computational innovation and biological application, we developed FishMamba-1 Hub (https://huggingface.co/spaces/lu1000001/FishMamba-Hub), a user-friendly web server powered by our trained models (Figure 7). The platform offers two core functionalities: 1, Sequence Encoder: Researchers can upload FASTA sequences to extract high-dimensional embeddings from the pre-trained FishMamba-1 backbone, enabling downstream tasks such as phylogeny construction or variant effect prediction. 2, Online Segmentation: A “drag-and-drop” interface allows users to input raw DNA sequences (up to 32k bp) and receive real-time, single-nucleotide resolution annotations of gene structures. The visualization module renders predicted exons, introns, and promoters as intuitive tracks (similar to Figure 4), democratizing access to advanced genomic AI for aquaculture researchers without coding expertise.

**Figure 7.**
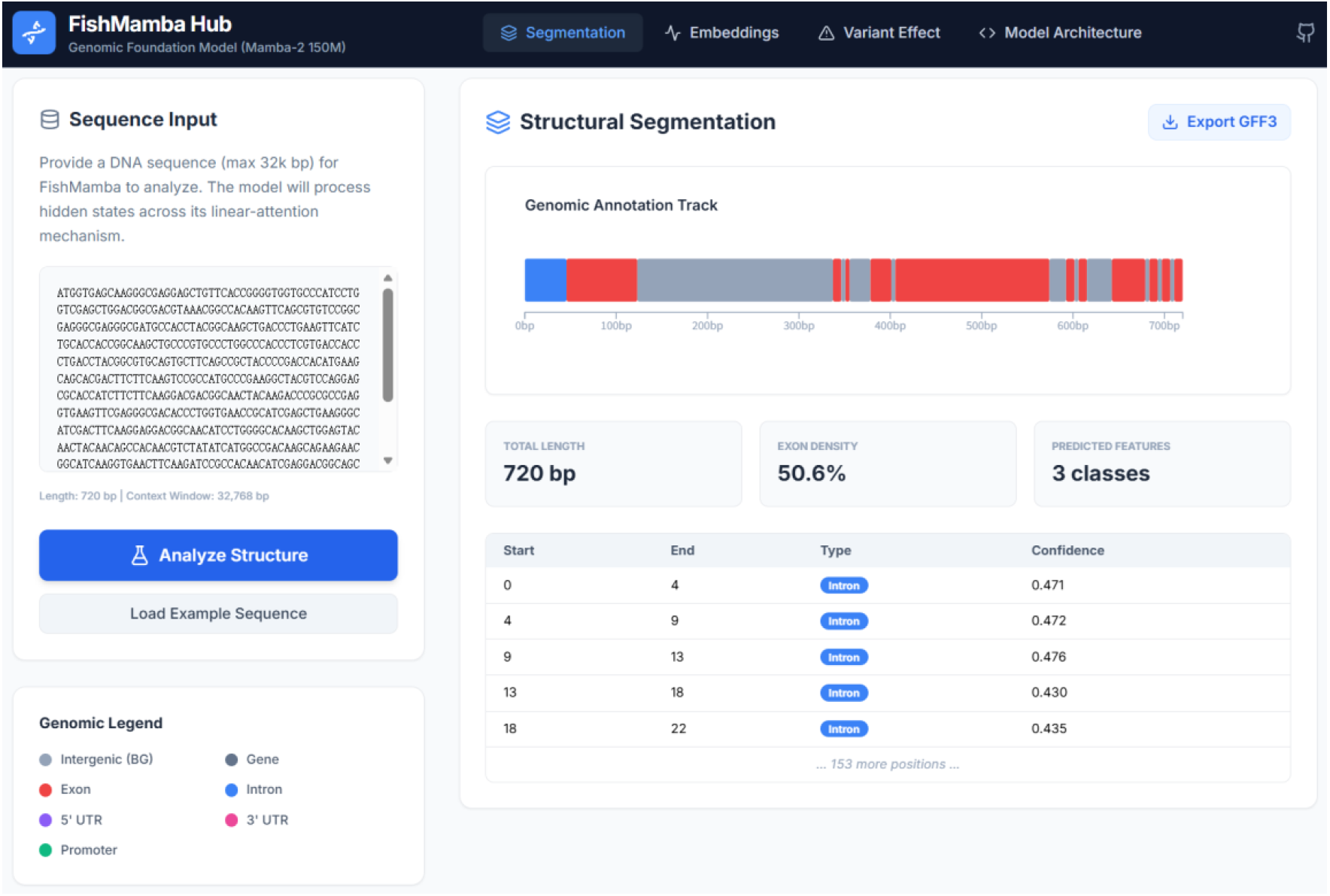
User interface of the FishMamba-1 Hub. The web-based tool allows researchers to upload or paste DNA sequences (up to 32k bp) for instant analysis. The interface displays predicted genomic structures (e.g., Exons, Promoters) through an intuitive color-coded track and a detailed data table. The platform supports GFF3 export, facilitating seamless integration into biological research workflows.

## 4. Discussion

### 4.1 Bridging the Context Gap in Polyploid Genomes

The genomic landscape of Cypriniformes is defined by its evolutionary complexity, characterized by multiple rounds of whole-genome duplication (WGD) and a consequent proliferation of repetitive elements (Xu et al., 2014; Yang et al., 2015). Decoding the “language” of such expanded genomes presents a computational paradox: biological signals are often separated by vast genomic distances, yet traditional Transformer-based models are restricted to short context windows (typically 4–6 kb) due to quadratic complexity (Nguyen et al., 2023). While recent benchmarks have debated the necessity of ultra-long contexts for general tasks (Feng et al., 2025), our results argue that for aquatic genomics, length matters. Cyprinid genomes are notorious for expanded introns driven by transposon insertions. By leveraging the Mamba-2 state-space architecture, FishMamba-1 achieves linear complexity, enabling a 32k context window. This 5-to-8-fold expansion allows the model to perceive the complete syntactic structure of expanded gene loci on standard academic hardware (e.g., NVIDIA A100), democratizing access to large-scale genomic AI.

### 4.2 The “False Positive” Paradox: Sequence Potential vs. Transcriptional Reality

A critical examination of our FishSegmenter results reveals a nuanced reality behind the so-called “false positives” in exon prediction. While traditional metrics might penalize these predictions (Figure 4), we argue they often reflect a limitation of the ground truth rather than the model. Current GFF3 annotations are largely derived from RNA-Seq data, which capture only the transcriptional reality of specific tissues at specific time points (Ji et al., 2021; Marand and Schmitz, 2022). These methods are known to sometimes under-estimate expression levels or miss genes due to genomic complexities and experimental conditions (Hirsch et al., 2015; Chisanga et al., 2022; Marand and Schmitz, 2022). In contrast, FishMamba-1 predicts biological potential directly from the DNA sequence. Consequently, many “false positives” likely represent unannotated exons, cryptic splicing events, or alternative isoforms that are biologically valid at the sequence level (containing correct AG/GT sites) but were transcriptionally silent or lowly expressed in the reference samples (Jaganathan et al., 2019; Koo and Eddy, 2019; Mehta et al., 2023). This suggests FishMamba-1 functions not just as an annotator, but as a generative discovery tool, capable of uncovering hidden coding potential in non-model organisms where transcriptomic data is sparse and annotations are often incomplete (Ji et al., 2021; Chisanga et al., 2022).

### 4.3 Global Semantic Understanding vs. Local Pattern Matching

Our comparative analysis with the CNN baseline uncovered a fascinating dichotomy. Specifically, the confusion matrix analysis (Figure 5) highlights the limitation of local receptive fields inherent to standard convolutional architectures (Kelley et al., 2018; Avsec et al., 2021). The CNN baseline’s inability to distinguish Intergenic regions from Introns (misclassifying 70% of the former) indicates a lack of awareness regarding “genic” versus “intergenic” states, a problem often attributed to the difficulty CNNs face in aggregating information over extremely long distances (Avsec et al., 2021). In contrast, FishMamba-1, leveraging its 32k context, maintains a persistent state representation that allows it to correctly assign non-coding sequences to their respective genomic compartments, solving a long-standing challenge in de novo annotation (Nguyen et al., 2023; de Almeida et al., 2025).

Conversely, the CNN baseline achieved a higher AUC on the specific task of splice site classification (Figure 6). This aligns with the inherent inductive bias of CNNs, which act as powerful “motif scanners” for local, translation-invariant patterns such as the canonical AG/GT splice sites (Jaganathan et al., 2019; Koo and Eddy, 2019). However, this local precision comes at a cost. FishMamba-1 prioritizes global semantic understanding. While it may miss isolated splice sites lacking broader contextual support, it excels at distinguishing broad genomic contexts (e.g., distinguishing Introns from Intergenic regions, Table 2) where long-range dependency is key (Nguyen et al., 2023). This trade-off highlights that while CNNs are excellent feature detectors, Foundation Models like FishMamba-1 are better genomic reasoners. Future work could explore hybrid architectures—combining the local sharpness of CNNs with the global reasoning of Mamba—to achieve the “best of both worlds” (Avsec et al., 2021; Schiff et al., 2024).

### 4.4 The Necessity of Clade-Specific Foundation Models

Finally, our work validates the “specialization hypothesis” in the aquatic domain. Just as AgroNT (Mendoza-Revilla et al., 2024) and PDLLMs (Liu et al., 2025a) established the need for plant-specific models to handle high heterozygosity, FishMamba-1 proves that generic human-centric models are insufficient for the unique evolutionary trajectories of fish. By pre-training on Cypri-24—a curated dataset covering ∼28.8 Gb of diverse cyprinid genomes— FishMamba-1 learned a generalized “fish syntax” shaped by sub-genome dominance and WGD. This specialization enables high-precision segmentation on “orphan” species not seen during training, providing a critical resource for molecular breeding in aquaculture.

### 4.5 Limitations and Future Directions

We acknowledge current limitations. While exon detection is robust, the identification of Untranslated Regions (UTRs) remains challenging (mIoU near zero), primarily due to the scarcity and incompleteness of UTR annotations in current non-model fish datasets rather than model incapacity; a challenge also noted in human segmentation models like SegmentNT (de Almeida et al., 2025). Furthermore, while FishSegmenter currently relies on token-level classification, recent breakthroughs highlight the efficacy of strict biological grammar enforcement. For instance, the Helixer model (Holst et al., 2025) achieves extraordinary precision by integrating a neural network backbone with a downstream HMM to resolve state transitions and reading frame phases at a biologically plausible level. In future iterations, we plan to augment the FishMamba-1 framework with a similar hybrid post-processing module and a phase-aware loss function. This will effectively bridge FishMamba-1’s linear-complexity global reasoning with exact, nucleotide-level boundary precision. Future iterations will benefit from integrating multi-modal data, such as ATAC-seq and RNA-seq, to refine regulatory element boundaries. Additionally, extending the context window to the 1M token scale could unlock the ability to model entire topologically associating domains (TADs).

## 5. Conclusion

FishMamba-1 demonstrates a critical transition from traditional homology-based annotations to sequence-driven foundational modeling in aquatic genomics. By validating the linear-complexity advantage of SSMs, we provide the community with a scalable, open-source framework that not only replicates known annotations but also uncovers the hidden coding landscape of polyploid genomes.

## Conflict of Interest

The authors declare no competing interests.

## Funding

This work was supported by the Jing-Jin-Ji Regional Integrated Environmental Improvement-National Science and Technology Major Project of Ministry of Ecology and Environment of China (No. 2025ZD1200802) and the National Natural Science Foundation of China (42330405, 32161160321, 32293252, and 32022009). This work was further supported by funding from the Hainan Provincial Key Research and Development Program ZDYF2025GXJS149.

## Data and Code Availability

The curated Cypri-24 genomic dataset, comprising 28.8 Gb of high-quality assemblies and standardized annotations used for pre-training and evaluation, is available via the project repository. The source code for FishMamba-1, including architectures, pre-training pipelines, and fine-tuning scripts, is publicly hosted on GitHub (https://github.com/lu1000001/FishMamba) under the MIT License. The pre-trained model weights and the fine-tuned FishSegmenter weights have been deposited in the Hugging Face Model Hub (https://huggingface.co/lu1000001/FishMamba). Furthermore, an interactive, web-based inference platform, FishMamba Hub, is accessible at https://huggingface.co/spaces/lu1000001/FishMamba-Hub, providing real-time, single-nucleotide resolution genome segmentation for the research community.

## Notes

https://github.com/lu1000001/FishMamba

https://huggingface.co/spaces/lu1000001/FishMamba-Hub

### Competing Interest Statement

The authors have declared no competing interest.

